# Interactome analysis of *C. elegans* synapses by TurboID-based proximity labeling

**DOI:** 10.1101/2021.04.01.438103

**Authors:** Murat Artan, Stephen Barratt, Sean M. Flynn, Farida Begum, Mark Skehel, Armel Nicolas, Mario de Bono

## Abstract

Proximity labelling provides a powerful *in vivo* tool to characterize the proteome of sub-cellular structures and the interactome of specific proteins. Using the highly active biotin ligase TurboID, we optimize a proximity labelling protocol for *C. elegans*. We use this protocol to characterise the proteomes of the worm’s gut, muscle, skin, and nervous system. We express TurboID exclusively in the pair of AFD neurons and show we can identify known and previously unknown proteins expressed selectively in AFD. We knock TurboID into the endogenous *elks-1* gene, which encodes a presynaptic active zone protein. We identify many known ELKS-1 interacting proteins as well as previously uncharacterised synaptic proteins. Versatile vectors, and the inherent advantage of *C. elegans* for biochemistry, make proximity labelling a valuable addition to the nematode’s armory.

**Teaser:** We optimize a TurboID proximity labeling protocol for C. elegans and use it to characterize tissue and synaptic proteomes

## Introduction

Characterizing the interactomes of specific proteins, and the proteome profiles of subcellular structures, cells and tissues, provides a powerful entry point to probe molecular function. Several methods designed to highlight protein-protein interactions (PPIs) have proven useful, including yeast-two-hybrid (Y2H), affinity purification and phage display (*1–5*). However, each method has limitations that can include high false positive rates, poor detection of transient or weak interactors, a low signal-to-noise ratio when detecting PPIs in specific cell types or subcellular compartments, artefacts created during tissue homogenization, and competing requirements for solubilizing proteins while keeping complexes intact (*1, 2*). Proximity-labeling methods overcome many of these limitations (*1, 6–8*), and have allowed the proteomes of subcellular compartments (*1*), and weak or transient PPIs to be characterized *in vivo* (*1*). Proximity labeling fuses a protein of interest to an enzyme domain that promiscuously tags proteins in its vicinity with a biochemical handle. This handle allows selective recovery of tagged proteins, which can then be identified by mass spectrometry (MS) (*1, 2, 7*). The enzyme domains most widely used for proximity-labeling in living cells and organisms are engineered variants of the *E. coli* biotin ligase BirA, e.g. BioID and TurboID, or of ascorbate peroxidase, e.g. APEX. *In vivo* applications in animals have, however, been relatively limited (*9, 10*), and required genetic modifications to alter cuticle permeability in the nematode (*11, 12*) and tissue dissection to increase H_2_O_2_ delivery to tissues (*13*) or pretreatment of live samples with detergent to increase biotin-phenol permeability in the fly (*14*).

*Caenorhabditis elegans* has proven a useful workhorse to investigate metazoan biology, offering powerful genetics, *in vivo* cell biology, an anatomy described at electron micrograph resolution, and a defined number of cells whose gene expression can be profiled at single cell resolution (*15–18*). The ability to probe protein complexes *in vivo* using proximity labeling would add a new dimension to studies in this animal. *C. elegans* offers advantages for proximity labeling. Fusion protein knock-ins can be generated easily, and functionally tested. Gram quantities of worms can be grown quickly and cheaply. Proximity labeling can be restricted to specific cell types, using appropriate promoters (*18–20*). Interacting proteins can be rapidly interrogated using CRISPR/Cas9-generated gene knockouts or knockdowns, e.g. using auxin-induced degradation (*21*).

*C. elegans* grows optimally at 15 – 25°C. BioID functions poorly at temperatures below 37°C (*22*), making it sub-optimal for use in the nematode. To date, there is no report applying BioID in *C. elegans*. However, BirA expressed in specific *C. elegans* tissues could biotinylate co-expressed proteins fused to an Avi tag, a 15-residue peptide substrate of BirA (*23*). The biotinylated Avi-tag fusion protein could then be affinity purified (AP) and co-immunoprecipitating proteins identified by MS (*24*). Reinke et al. (*11, 12*) expressed cytosolic or nuclear APEX in various *C. elegans* tissues and characterized the corresponding proteomes. APEX has not, however, been used to study PPIs in worms. The APEX peroxidase is promiscuous, labeling proteins far from the APEX-tagged protein and leading to specificity problems (*25*). Two other studies characterized tissue proteomes in *C. elegans* by expressing mutant phenylalanyl tRNA synthetase (MuPheRS) (*26, 27*). MuPheRS can charge Phe tRNAs with azido-phenylalanine, so that this non-canonical amino acid can be incorporated into proteins during translation. Click chemistry permits the azido group to be derivatized and labeled proteins affinity purified for mass spectrometry. These studies highlighted the proteome of various tissues, including neurons. MuPheRS cannot, however, be used to study PPIs.

More catalytically active variants of BioID, called TurboID and miniTurbo, have recently been developed using directed evolution (*28*). Expressing either variant in *C. elegans* results in robust biotinylation signals in the intestine, with the strongest signals generated by TurboID (*28*). However, the authors did not publish interactome or proteomic data for *C. elegans.* A recent study used TurboID to characterize the interactome of the microtubule-binding protein patronin, but ectopically overexpressed this fusion protein in the largest tissue of the animal, the gut, and did not seek to optimise the protocol (*29*).

Here, we optimize a protocol for TurboID-based proximity-labeling in *C. elegans*. We express TurboID in different *C. elegans* tissues and in the pair of AFD thermosensory neurons. We also knock TurboID into *elks-1*, which encodes a pre-synaptic active zone protein. We characterize tissue proteomes and highlight tissue-specific proteins. Targeting TurboID to AFD neurons highlights the AFD-specific proteins GCY-8, TTX-1 and GCY-18, and previously uncharacterized proteins that we show are selectively expressed in AFD. The ELKS-1-TurboID samples are enriched in the synaptic proteins UNC-10/RIM, SYD-1/SYDE1, SYD-2/liprin-alpha, SAD-1/BRSK1, CLA-1/CLArinet, C16E9.2/Sentryn, RIM binding protein RIMB-1, and previously uncharacterized proteins, which we show localize at synapses. Our results indicate TurboID-mediated proximity labeling can effectively identify the proteome of a pair of neurons and bona fide interactors of a synaptic protein in *C. elegans in vivo*.

## Results

### Optimizing proximity labeling using TurboID in *C. elegans*

To examine the extent to which TurboID biotinylates proteins in *C. elegans*, we generated transgenic animals expressing a TurboID-mNeongreen (TbID-mNG) fusion protein in various tissues (Fig. 1A). We compared protein biotinylation levels in age synchronized young adults expressing the neuronal *rab-3p::TbID-mNG* transgene with wild type controls. In the absence of exogenously added biotin, transgenic animals showed only a slight increase in biotinylated proteins compared to controls. Adding exogenous biotin increased protein biotinylation specifically in animals expressing the *rab-3p::TbID-mNG* transgene (Fig. 1B). To optimize biotin availability to worm tissues, we treated animals with exogenous biotin for different time intervals, using biotinylation in neurons as a readout. A two-hour incubation was sufficient to achieve robust protein labeling (Fig. 1C). To identify an optimum biotin concentration for protein labeling, we treated animals with varying concentrations of biotin for 2 hours. We observed a substantial increase in biotinylation in worms treated with 1 mM biotin but higher biotin concentrations did not appear to further increase biotinylation (Fig. 1D). We also used the *E. coli* biotin auxotrophic strain MG1655 as a food source for worms instead of standard OP50 (*30*), to minimize the free biotin available prior to addition of exogenous biotin, allowing for tighter control of the time window during which promiscuous biotinylation occurs.

**Fig. 1.**
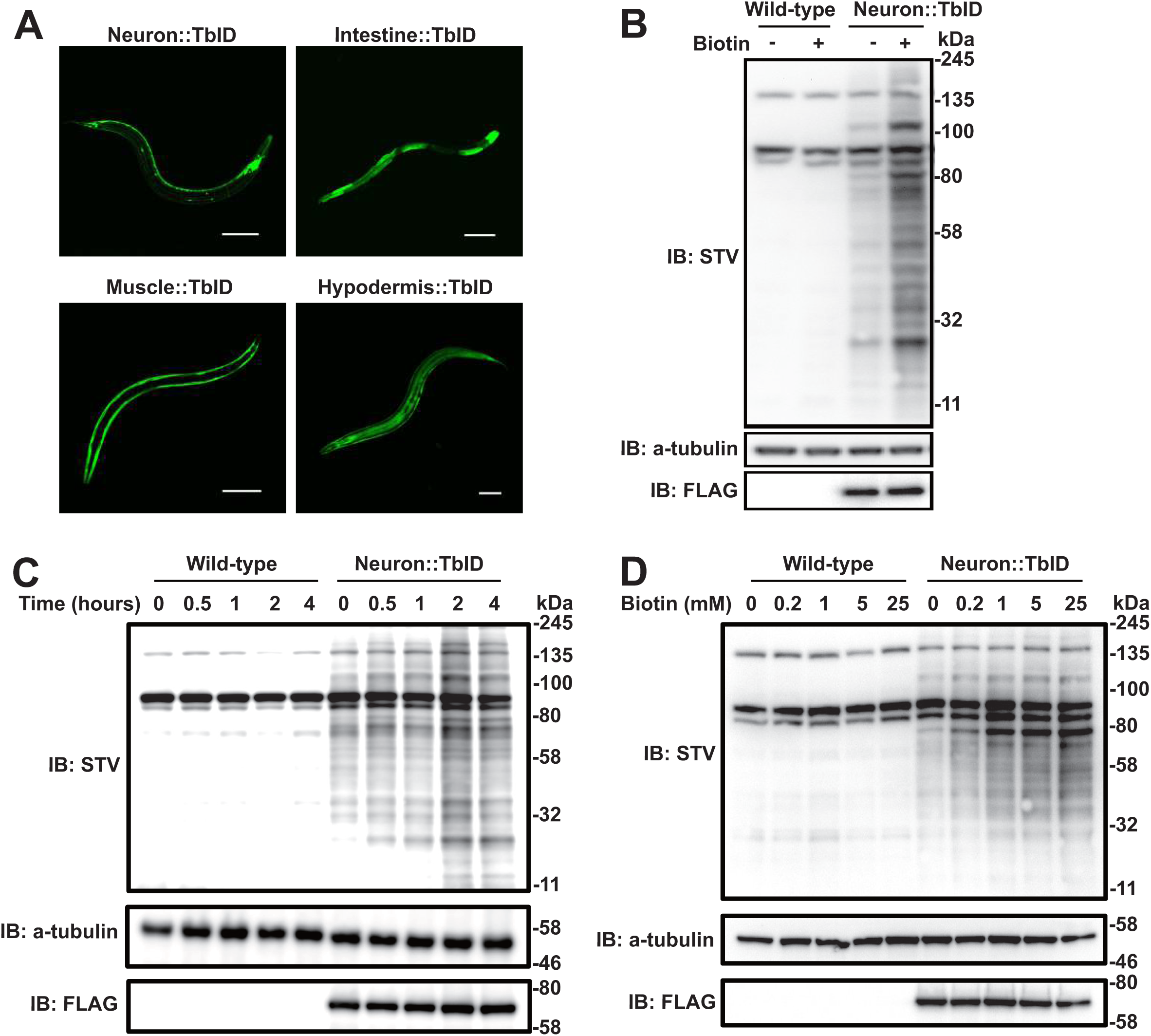
Optimizing TurboID-mediated proximity labeling in *C. elegans*. (**A**) Transgenic *C. elegans* expressing a free TurboID::mNeongreen::3xFLAG fusion protein selectively in neurons (*rab-3p*), intestine (*ges-1p*), body wall muscle (*myo-3p*) and hypodermis (*dpy-7p*). (**B**) Western blot analysis of biotinylation by TurboID in the nervous system (*rab-3p::TbID*) in the presence or absence of excess exogenous biotin. (**C** and **D**) Analyses of how labeling time (**C**) and exogenous biotin concentration (**D**) alters biotinylation in animals expressing TurboID pan-neuronally. Wild-type worms are shown as controls. Scale bars: 100 µm.

### Robust biotinylation in different *C. elegans* tissues expressing TurboID

Like pan-neuronal expression, intestinal, hypodermal and muscle-specific expression of TurboID-mNG conferred robust biotinylation activity (Fig. 2A), indicating that TurboID is functional in all major *C. elegans* tissues. We next optimized a protocol to extract and affinity purify biotinylated proteins from *C. elegans* (Fig. S1A-C). A significant advantage of PL compared to IP is that extracts can be collected under strong denaturing conditions (1% SDS, 2M urea), since the biotin tag is covalently attached. This allowed us to achieve >95 % solubilization of proteins (Fig. S1A–C) Methods). We achieved efficient protein extraction using a cryomill, and processed from 5 g to < 500 mgs of worms; using 500 mg of worms was typically sufficient to obtain enough material for MS. Age synchronized young adult worms were incubated with 1 mM of biotin for 2 hours at room temperature and immediately frozen after three washes with M9 buffer. Total protein lysate was extracted from powdered worm samples, and passed through desalting columns to eliminate free biotin. Removing free biotin was key for successful affinity purification. The flow through was incubated with streptavidin Dynabeads to affinity purify biotinylated proteins (see Methods). Western blot analysis showed capture of the majority of biotinylated proteins from the whole lysate using our affinity purification protocol (Fig. S2A). After extensive washing and elution, we fractionated the affinity-purified biotinylated proteins by SDS-PAGE, visualized the proteins by Coomassie staining, and analysed gel slices by LC-MS/MS analysis (Fig. S2B). Using a threshold of at least two unique peptides, we cumulatively identified >4000 proteins expressed in one or more tissues (Fig. 2B and C and Table S1). Using tissue enrichment analysis (*31*), we looked for over-represented annotations in the lists of proteins we identified as unique to each tissue. As expected, terms associated with the targeted tissue were over-represented in each case, confirming the validity of our method in large tissues (Fig. S3A). In addition, we observed a strong correlation between the two replicates at the level of spectral counts (Fig. S3B). Consistent with previous findings, we identified most proteins in the intestine, followed by the hypodermis and the muscle cells (Fig. 2B), (*12*). Over 2000 proteins were enriched at least two-fold in one tissue over other tissues (Fig. 2B), and 1274 proteins were detected in only one tissue (Fig. 2C). Proteins identified by MS/MS when TurboID was targeted to neurons included broadly expressed neuronal proteins, such as CLA-1, UNC-31, and proteins expressed in subsets of neurons, such as the GABA vesicular transporter UNC-25, expressed in 26 neurons, and OSM-10 which is expressed in 4 pairs of neurons (Fig. 2D). Muscle-specific samples were enriched in CPNA-1, CPNA-2, PQN-22 and F21C10.7 (Fig. 2D); intestine-specific samples were enriched in ACOX-2, CGR-1, C49A9.9 and C49A9.3 (Fig. 2D), and hypodermis-specific samples included F17A9.5, PAH-1, R05H10.1 and CPN-2 (Fig. 2D). In addition, we identified tissue-specific enrichment for many proteins whose expression was previously uncharacterized (Table S1 and Fig. 2E). For some of these proteins we validated the tissue-selective expression highlighted by our MS/MS data by making transgenic reporters. The reporters confirmed the expression profile predicted by MS/MS, validating the method’s specificity (Fig. 2F).

**Fig. 2.**
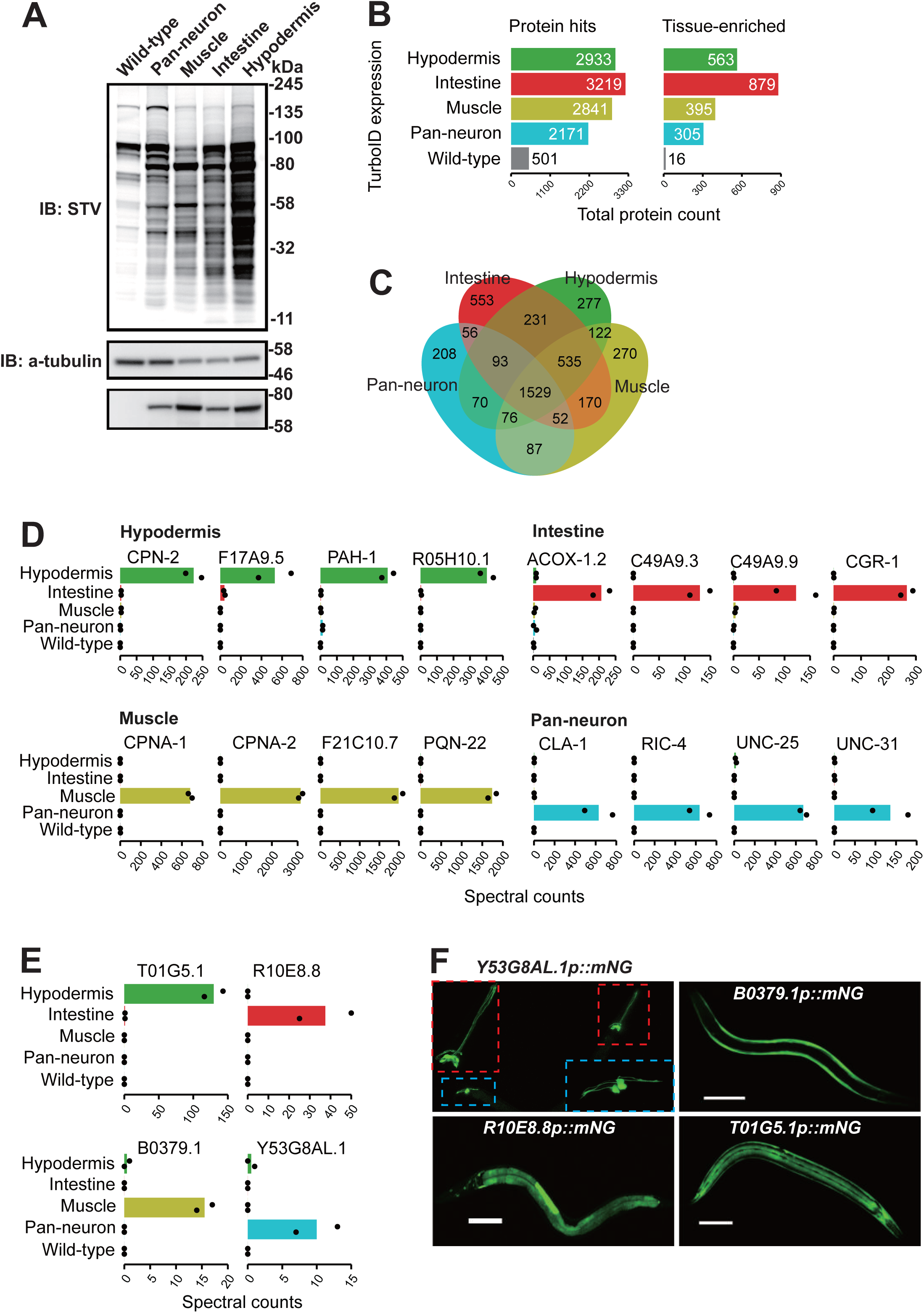
Proteomic analysis of major *C. elegans* tissues. (**A**) Western blot of lysates obtained from animals expressing free TurboID specifically in neurons, muscle, intestine and hypodermis. (**B**) Total, tissue-enriched and tissue-specific proteins detected in various tissues by mass spectrometry. Tissue-enrichment is defined as proteins with twofold or greater mean spectral counts compared to the second-ranked sample. Tissue-specific proteins were uniquely identified in the corresponding tissue. (**C**) Venn diagram showing the distribution of protein hits between samples. (**D**) Mean spectral count of representative proteins in various tissues obtained via mass spectrometry. (**E** and **F**) Mean spectral counts of the proteins Y53G8AL.1, B0379.1, T01G5.1 and R10E8.8 encoded by previously uncharacterized genes predicted by our TurboID experiments (E), and confocal microscopy images of worms expressing mNeongreen from the promoters of these genes (F). Scale bars: 100 µm.

### TurboID can identify proteins expressed in only a pair of *C. elegans* neurons

The *C. elegans* nervous system includes 118 classes of neurons, with most classes consisting of a single pair of neurons that form left/right homologs (*16*). We asked whether our protocol had sufficient sensitivity to characterize proteins expressed in a single pair of neurons. To test this, we transgenically expressed free TurboID specifically in the AFD pair of ciliated sensory neurons, using the *gcy-8* promoter (Fig. 3A). Western blot analysis of extracts from these animals revealed a biotinylation signal significantly higher than that in extracts from wild-type controls (Fig. 3B). Correlation plots of mass spectrometry data obtained for affinity-purified extracts made from animals expressing the *gcy-8p::TurboID::mNG* transgene showed reproducible results between replicates (Fig. S4A). As expected, these samples were enriched for proteins specifically or selectively expressed in AFD compared to similarly processed extracts from non-transgenic controls, or from animals expressing a *rab-3p::TurboID::mNG* transgene (Fig. 3C and S4B). Enriched proteins included the transmembrane guanylate cyclases GCY-8 and GCY-18, and the homeobox transcription factor TTX-1 (Fig. 3D and S4B). Our MS data also identified other proteins enriched in the AFD-specific TurboID samples compared to the pan-neuronal TurboID controls (Table S1, Fig. 3E and S4C). To examine if these proteins were selectively expressed in AFD neurons, we generated transgenic reporter lines (Fig. 3F). *nex-4* and F37A4.6 were expressed specifically in AFD, albeit F37A4.6 at low levels; T06G6.3 was expressed in a small subset of neurons that included AFD (Fig. 3F). Our list of AFD-enriched proteins, identified by mass spectrometry, was consistent with AFD-specific gene expression identified by RNA Seq (*32*). These data suggest that TurboID is a reliable tool to map the proteome of specific neurons in *C. elegans*.

**Fig. 3.**
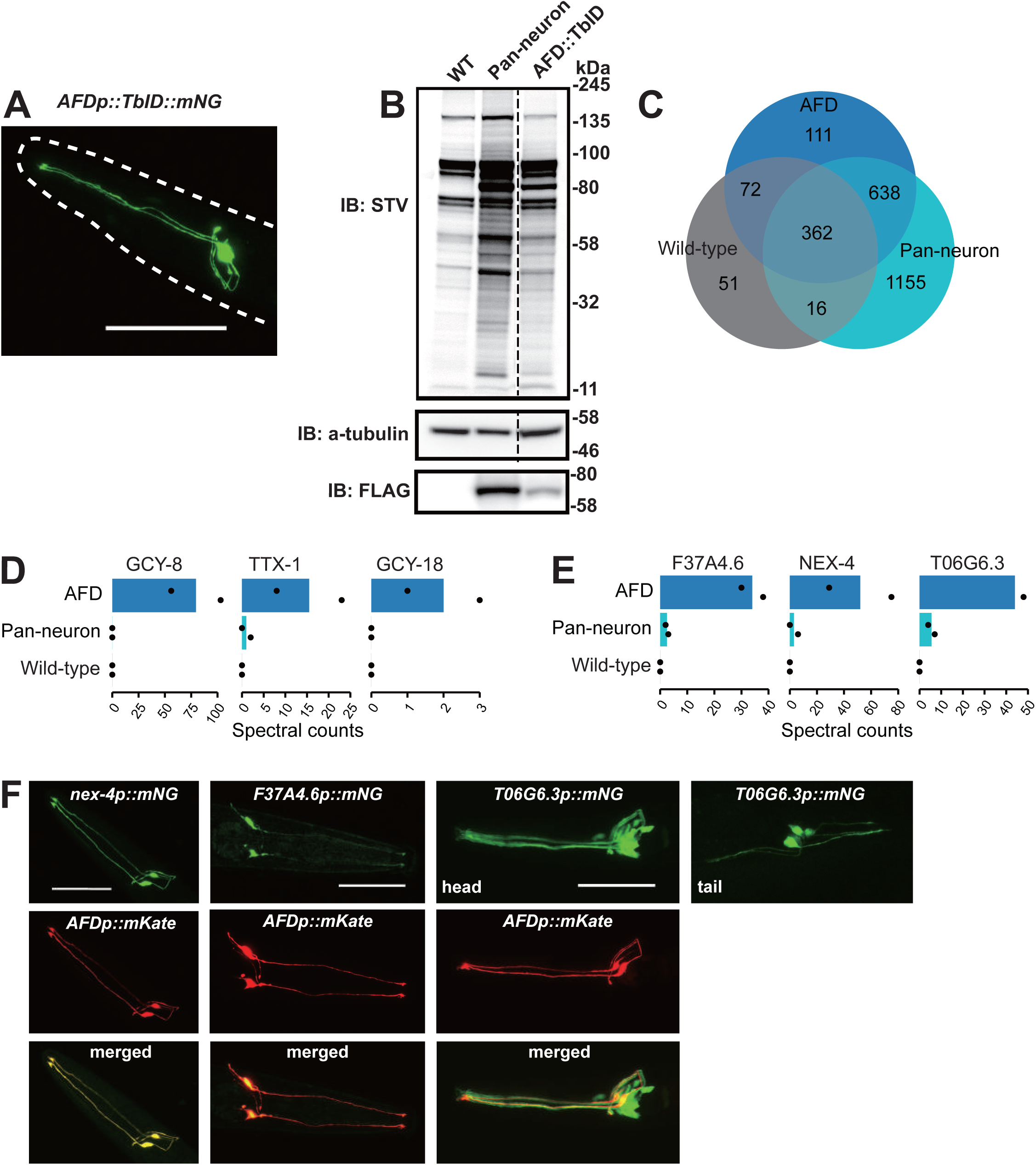
Proteomic analysis of the AFD sensory neuron pair. (**A**) Confocal microscopy image of *C. elegans* expressing TurboID::mNeongreen fusion protein specifically in the AFD neurons. (**B**) Western blot analysis of worm lysates obtained from samples expressing free TurboID in AFD neurons, all neurons or none (WT). (**C**) Venn diagram showing the distribution of protein hits between samples. (**D**) The AFD markers GCY-8, TTX-1 and GCY-18 are highly enriched in AFD::TurboID samples. (**E**) NEX-4, F37A4.6 and T06G6.3 proteins were highly enriched in AFD::TurboID samples. (**F**) Confocal microscopy images of *C. elegans* expressing *nex-4p::mNeongreen*, *F37A4.6p::mNeongreen* and *T06G6.3p::mNeongreen* transgenes. A *gcy-8p::mKate* transgene was used as a fiducial marker to identify AFD neurons. Scale bars: 50 µm.

### Identifying the interactome of a pre-synaptic protein by TurboID

We next asked if TurboID can highlight the interactome of a specific *C. elegans* protein expressed at endogenous levels. We focused on the synaptic protein ELKS-1, an ortholog of human ERC2 (ELKS/RAB6-interacting/CAST family member 2). ELKS-1 is expressed throughout the nervous system and localizes to the pre-synaptic active zone, in proximity to other pre-synaptic proteins such as α-liprin and RIM (*33*). To have endogenous levels of expression, we knocked *TurboID::mNeongreen* into the *elks-1* locus using Crispr/Cas9-mediated genome editing. As expected, ELKS-1::TurboID::mNeongreen localized to the nerve ring and other regions rich in synapses (Fig. 4A). Western blot analysis of extracts from ELKS-1::TbID animals did not show an increased biotinylation signal compared to wild-type controls (Fig. 4B). However, mass spectrometry analysis of streptavidin-purified proteins from this knock-in strain revealed enrichment of known synaptic proteins including UNC-10/RIM, SYD-1/SYDE1, SYD-2/α-liprin, SAD-1/BRSK1, CLA-1/CLArinet, C16E9.2/Sentryn and the RIM binding protein RIMB-1 (Figure 4D). We measured enrichment by comparing mass spectrometry data obtained for control extracts processed in parallel from wild type and from transgenic animals expressing free TurboID-mNG throughout the nervous system, *rab-3p::TurboID-mNG* (Fig. 4C and S4E) or both nervous system and other tissues (Fig. S4G). Between-replicate correlation plots revealed reproducible results between experimental repeats (Fig. S4D). We also identified several previously uncharacterized proteins as enriched in ELKS-1::TurboID samples (Table S1, Fig. 4E and S4F and G). We expressed mNeongreen translational fusion transgenes for several of these proteins exclusively in AFD neuron pair, and showed that they co-localized with ELKS-1::mScarlet at pre-synaptic active zone where AFD synapses with AIY interneurons (*16, 34*) (Fig. 4E and F; highlighted with red dots). Together, our findings show that TurboID-mediated proximity labeling is an effective method to reveal protein interactors in *C. elegans* at endogenous levels with specificity and sensitivity.

**Fig. 4.**
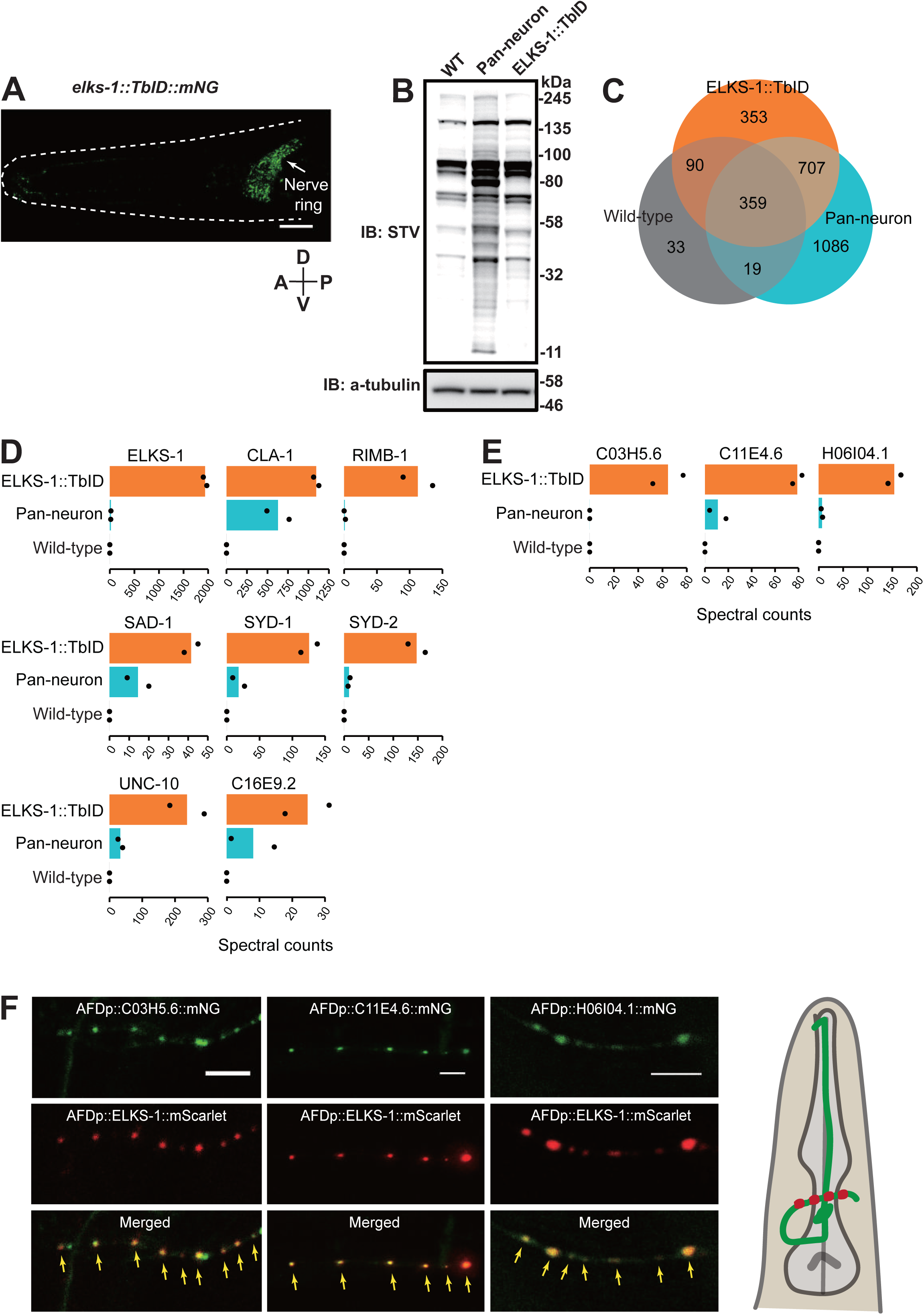
Interactome analysis of the pre-synaptic active zone protein ELKS-1. (**A**) Confocal microscopy image of *C. elegans* harboring an *elks-1::TurboID::mNeongreen* knock-in allele generated by CRISPR/Cas9-mediated genome editing. The nerve ring, where many synapses are located, is marked with a white arrow. (**B**) Western blot analysis of lysates obtained from animals expressing the ELKS-1::TurboID::mNeongreen fusion protein. (**C**) Venn diagram showing the distribution of protein hits between samples. (**D**) The synaptic proteins ELKS-1/ELKS, UNC-10/RIM, SYD-1/SYDE1, SYD-2/liprin-alpha, SAD-1/BRSK1, CLA-1/CLArinet, C16E9.2/Sentryn and RIM binding protein RIMB-1 were highly enriched in ELKS-1::TurboID samples. (**E**) C03H5.6, C11E4.6 and H06I04.1 were highly enriched in ELKS-1::TurboID samples. (**F**) Confocal microscopy images of transgenic *C. elegans* expressing *C03H5.6::mNeongreen*, *C11E4.6::mNeongreen* and *H06I04.1::mNeongreen* translational fusions in the AFD neuron pair. *elks-1::mScarlet,* used as a control to mark AFD synapses, confirms these proteins are synaptic. Scale bars: 10 µm (**A**) and 5 µm (**F**).

## Discussion

We optimize the recently developed TurboID-based proximity-dependent protein labeling approach for *C. elegans,* and show that it permits single neuron proteomics and characterization of the interactome of a synaptic protein expressed at endogenous levels. We create reagents that allow TurboID to be applied to different *C. elegans* tissues and identify > 4000 proteins expressed in at least one tissue. We were able to identify proteins exclusively expressed in the AFD neurons, and previously uncharacterized proteins that are synaptically localized in the vicinity of ELKS-1.

Proximity labeling methods like TurboID label interactors throughout the life of the fusion protein. In *C. elegans*, many transgenic experiments use multi-copy arrays that typically overexpress the protein of interest. This can lead to inappropriate protein localization, expanding the list of interactors identified by proximity labeling. Expressing bait proteins at physiological levels, and confirming appropriate sub-cellular localization of the fusion protein, may be important to minimize such confounds. Expressing ELKS-1::TurboID at physiological levels, using knock-in strains, identified many proteins predicted to be enriched in the vicinity of ELKS-1. These include known interactors of ELKS-1 in the pre-synaptic active zone of *C. elegans* neurons. We characterize three proteins, C11E4.6, C03H5.6 and H06I04.1, enriched in proteins biotinylated by ELKS-1::TurboID further and confirm they are synaptically localized. C11E4.6 is predicted to be an 1170 residue protein orthologous to human ANKS1A and ANKS1B (ankyrin repeat and sterile alpha motif domain-containing). Human ANKS1B, also known as AIDA-1, is a risk locus for autism and neurodevelopmental defects (*35*), and is enriched at post-synaptic densities, where it binds N-methyl-D-aspartate receptors (NMDA) and the adapter protein PSD-95 (*36*). Our data are consistent with this family of proteins also being enriched pre-synaptically. Previous studies have reported co-localization of AIDA-1 with the pre-synaptic excitatory marker VGLUT1 in mossy fibre endings onto CA3 neurons (*37*), and some pre-synaptic proteins co-immunoprecipitate with AIDA-1 including the CLA-1 homolog Bassoon (*35*). C03H5.6 is predicted to encode a 349 residue protein that contains BTB/POZ domains; little is known about this protein’s function. H06I04.1 is predicted to encode four isoforms ranging from 231 to 428 residues and contains five coiled-coil domains. Similar to C03H5.6, little is known about this protein’s function. Transgenic animals overexpressing C11E4.6 exhibit an uncoordinated phenotype, supporting the idea that it is a synaptic protein. It would be interesting to investigate whether C11E4.6, C03H5.6 or H06I04.1 have a role in synaptic transmission. Another protein that our TurboID data suggest is proximal to ELKS-1 is C16E9.2, the *C. elegans* ortholog of human KIAA0930. C16E9.2 was recently given the name sentryn (STRN-1). STRN-1 together with the SAD kinase and liprin-α promote dense-core vesicle (DCV) pausing at pre-synaptic regions (*38*) and optimize localization of synaptic vesicles at the active zone (*39*).

An important variable for TurboID protocols is the delivery of biotin or biotin derivatives. In cultured mammalian cells expressing TurboID or miniTurbo, adding biotin for as little as 10 minutes yields robust biotinylation (*28*). In *C. elegans*, adding biotin to plates seeded with bacteria (*E. coli* OP50 or NA22 strains) is sufficient to promote TurboID labeling, but long incubation times are required to achieve detectable protein biotinylation (*28*; our data). We find that a 2-hour incubation with exogenous biotin is sufficient for strong TurboID signals in worm neurons (Fig. 1C). Since in the lab *C. elegans* is typically grown on *E. coli* as a food source, feeding biotin-auxotrophic *E. coli* to worms provides some control over the start of biotinylation by TurboID. For time-sensitive experiments, it may be possible to soak worms in a buffer containing biotin for shorter periods, e.g. 30 or 60 minutes and still achieve robust protein biotinylation, although further analysis using MS is required to confirm this.

Endogenously biotinylated proteins dominate the biotinylated proteome in TurboID expressing lines, as seen by both Western blot (Fig. S2A) and MS analysis (data not shown). Four carboxylases; PCCA-1/PCCA (propionyl coenzyme A carboxylase alpha subunit), PYC-1/PC (pyruvate carboxylase), MCCC-1/MCCC (methylcronotoyl coenzyme A carboxylase), POD-2/ACACA (acetyl coenzyme A carboxylase) were the most abundantly detected proteins in our MS experiments in every condition tested, including controls. Depending on how broadly and strongly we expressed TurboID in *C. elegans* tissues, these carboxylases accounted for 8% – 85% of peptides identified in MS analysis. We aim to generate transgenic animals that have peptide tags knocked into each of these four genes (*pcca-1*, *pyc-1*, *mccc-1* and *pod-2*), using CRISPR/Cas9. This would allow depletion of these endogenously biotinylated proteins by affinity purification, and is likely to increase the sensitivity of proximity labeling substantially, particularly when the TurboID-tagged protein is expressed in a small number of cells or at low levels.

In summary, our findings show that TurboID works well in a wide array of contexts in the worm. TurboID will be a reliable and useful tool for the *C. elegans* community to map the proteomes of specific cells and subcellular structures, and to characterize the interactomes of proteins of interest.

## Materials and methods

### Strains

Worms were grown at room temperature (22°C) on nematode growth medium (NGM) plates seeded with the biotin auxotroph *E. coli* MG1655. *C. elegans* husbandry otherwise followed standard laboratory culture conditions (*40*).

*C. elegans* strains used in this study include: N2, the wild-type Bristol strain

AX7647 *dbIs37[rab-3p::mNeongreen::3XFLAG]*

AX7526 *dbIs24[rab-3p::CeTurboID::mNeongreen::3XFLAG]*

AX7542 *dbIs25[ges-1p::CeTurboID::mNeongreen::3XFLAG]*

AX7578 *dbIs28[myo-3p::CeTurboID::mNeongreen::3XFLAG]*

AX7623 *dbIs33[dpy-7p::CeTurboID::mNeongreen::3XFLAG]*

AX7606 *dbIs32[gcy-8p::CeTurboID::mNeongreen::3XFLAG]*

AX7917 *dbEx1234[T01G5.1p::mNeongreen::3XFLAG; ccRFP]*

AX7922 *dbEx1241[B0379.1p::mNeongreen::3XFLAG; ccRFP]*

AX7928 *dbEx1247[Y53G8AL.1p::mNeongreen::3XFLAG; ccRFP]*

AX8059 *dbEx1266[R10E8.8p::mNeongreen::3XFLAG]*

AX7935 *dbEx1251[nex-4p::mNeongreen::3XFLAG; ccRFP]; dbEx1253[gcy-8p:: mKate; ccRFP]*

AX7937 *dbEx1255[F37A4.6p::mNeongreen::3XFLAG; lin-44p::GFP]; dbEx1253[gcy-8p:: mKate; ccRFP]*

AX7939 *dbEx1256[T06G6.3p::mNeongreen::3XFLAG]; dbEx1253[gcy-8p:: mKate; ccRFP]*

AX8110 *dbEx1296[gcy-8p::C03H5.6::mNeongreen::3XFLAG; gcy-8p::elks-1 cDNA::mScarlet; lin-44p::gfp]*

AX8111 *dbEx1297[gcy-8p::C11E4.6::mNeongreen::3XFLAG; gcy-8p::elks-1 cDNA::mScarlet; lin-44p::gfp]*

AX8117 *dbEx1300[gcy-8p::H06I04.1 cDNA isoform 1a::mNeongreen::3XFLAG; gcy-8p::elks-1 cDNA::mScarlet; lin-44p::gfp]*

PHX1710 *elks-1(syb1710)*. The *syb1710* allele is a knock-in that expresses ELKS-1 tagged C-terminally with CeTurboID::mNeongreen::3XFLAG.

### Molecular biology

#### TurboID cloning

TurboID codon-optimized for *C. elegans* (*41*) was synthesized by IDT (Integrated DNA Technologies Inc, Coralville, IA, USA) and includes an N-terminal Gly-Ser rich linker, a c-Myc tag C-terminally, and two artificial introns (N linker::CeTurboID::c-Myc).

*C. elegans* codon-optimized mNeongreen::3XFLAG tag was cloned using PCR from DG398 pEntryslot2_mNeongreen::3XFLAG::stop, a Multisite Gateway (ThermoFisher) pEntry vector which was a gift from Dr. Dominique Glauser (University of Fribourg).

DNA encoding N linker::CeTurboID::c-Myc and mNeongreen::3XFLAG were stitched together by fusion PCR using primer sequences that encoded a short Gly-Ser linker. The resulting PCR product (N linker::CeTurboID::c-Myc::mNeongreen::3XFLAG) was cloned into position 2 of a Multisite Gateway pDONR vector (ThermoFisher).

#### Transcriptional reporters

Promoters for B0379.1 (∼0.9 kbp), T01G5.1 (∼2.1 kbp), Y53G8AL.1 (∼2.5 kbp), nex-4 (∼1.6 kbp), F37A4.6 (∼2.5 kbp), T06G6.3 (∼2.2 kbp) and R10E8.8 (∼1.6 kbp) were amplified from *C. elegans* genomic DNA by PCR and cloned into position 1 of a Multisite Gateway Donor vector (ThermoFisher). The resulting pEntryslot1 vectors containing these promoters were each mixed with the dg398 pEntryslot2_mNeongreen::3XFLAG::stop vector, a pEntryslot3 vector containing the *let-858* or *tbb-2* 3’UTR and pDEST (ThermoFisher) in an LR reaction. The resulting Expression vectors were injected into the gonad of day 1 adult N2 worms at a concentration of 25 ng/uL.

#### Cloning uncharacterized ELKS-1 interactors

ORFs for *C03H5.6* (∼2 kbp) and *C11E4.6* (∼6 kbp) were amplified from *C. elegans* genomic DNA by PCR and cloned into a position 2 Gateway donor vector. cDNA for *H06I04.1* isoform a was synthesized by IDT (Integrated DNA Technologies Inc, Coralville, IA, USA) and cloned into a position 2 Gateway donor vector. The resulting pEntryslot2 vectors were mixed with dg397 pEntryslot3_mNeongreen::3XFLAG::stop::unc-54 3’UTR entry vector and a pDEST vector in an LR reaction. *elks-1* cDNA (∼2.5 kbp) was PCR-amplified from a *C. elegans* cDNA library and cloned into a position 2 Gateway donor vector, mixed with *gcy-8* promoter (position 1) and *mScarlet* (position 3) and a pDEST vector in an LR reaction. Resulting expression vectors were injected into the gonad of day 1 adult N2 worms at a concentration of 25 ng/uL.

#### Primers used for cloning

***myo-3* promoter (∼2.6 kb)**

F-myo-3p-Entry1- GGGGACAACTTTGTATAGAAAAGTTGTGTAGGCAATCAGTCAAACCGAATAAAA

R-myo-3p-Entry1-GGGGACTGCTTTTTTGTACAAACTTGTTCTAGATGGATCTAGTGGTCGTGGGTT

***dpy-7* promoter (∼380 bp)**

F-dpy-7p-Entry1-GGGGACAACTTTGTATAGAAAAGTTGAATCTCATTCCACGATTTCTCGCAACACA T

R-dpy-7p-Entry1-GGGGACTGCTTTTTTGTACAAACTTGTTATCTGGAACAAAATGTAAGAATATTCTT A

***gcy-8* promoter (∼940 bp)**

F-gcy-8p-Entry1- GGGGACAACTTTGTATAGAAAAGTTGGGTTCAACAAGGGTATTGTATTGCAATCA GTG

R-gcy-8p-Entry1- GGGGACTGCTTTTTTGTACAAACTTGTTTGATGTGGAAAAGGTAGAATCGAAAAT CC

***T01G5.1* promoter (∼2.1 kbp)**

F-T01G5.1p-Entry1- GGGGACAACTTTGTATAGAAAAGTTGCACCTGAACACAACATTTTCTG

R-T01G5.1p-Entry1- GGGGACTGCTTTTTTGTACAAACTTGGACATGAAATTGTATCTGAAAGC

***Y53G8AL.1* promoter (∼2.5 kbp)**

F-Y53G8AL.1p-Entry1- GGGGACAACTTTGTATAGAAAAGTTGGTAGTTATGGAAAAGCAACGTCGGAG R-Y53G8AL.1p-Entry1-

GGGGACTGCTTTTTTGTACAAACTTGTCTAAAATAGCATTGGTTCTGAAACTTTG

***B0379.1* promoter (∼0.9 kbp)**

F-B0379.1p-Entry1- GGGGACAACTTTGTATAGAAAAGTTGGAAACAATATTATTTTTGTTTCACAG

R-B0379.1p-Entry1- GGGGACTGCTTTTTTGTACAAACTTGGTTGTTGTTGTCTCGATGGAAAAG

***nex-4* promoter (∼1.6 kbp)**

F-nex-4p-Entry1- GGGGACAACTTTGTATAGAAAAGTTGCCTGAGAATTACTGAAGTTTAAGC

R-nex-4p-Entry1- GGGGACTGCTTTTTTGTACAAACTTGCTCGAGTTACTTCAATGCTCACG

***F37A4.6* promoter (∼2.5 kbp)**

F-F37A4.6p-Entry1- GGGGACAACTTTGTATAGAAAAGTTGGATTCAGAAAATTCAGAAAGGCATTC

R-F37A4.6p-Entry1- GGGGACTGCTTTTTTGTACAAACTTGCATTATAGGAAGACTGAGATTCCAAGC

***T06G6.3* promoter (∼2.2 kbp)**

F-T06G6.3p-Entry1- GGGGACAACTTTGTATAGAAAAGTTGCATTTTTTGCTCTAAAGGTGGAATAG

R-T06G6.3p-Entry1-GGGGACTGCTTTTTTGTACAAACTTGCTTTGAAAAAAGTTCAGAGTAGTAGAG

***C03H5.6* genomic DNA (∼2 kbp)**

F-C03H5.6-Entry2- GGGGACAAGTTTGTACAAAAAAGCAGGCTtttcagaaaaATGTTAGAATGTATACATCCAACAT

R-C03H5.6-Entry2- GGGGACCACTTTGTACAAGAAAGCTGGGTATGTACGATTCATCAAACCACCTAC

***C11E4.6* genomic DNA (∼6 kbp)**

F-C11E4.6-Entry2- GGGGACAAGTTTGTACAAAAAAGCAGGCTtttcagaaaaATGAGCCTCAAAGACTTTGTCATATC

R-C11E4.6-Entry2- GGGGACCACTTTGTACAAGAAAGCTGGGTAATTTCTTTGGTTCTCAGTAGTTTGC TG

***H06I04.1* cDNA isoform 1a (∼1.5 kbp)**

F-H06I04.1-Entry2- GGGGACAAGTTTGTACAAAAAAGCAGGCTtttcagaaaaATGAGCAAAGAAACTGGAAAAATGGCGG

R- H06I04.1-Entry2- GGGGACCACTTTGTACAAGAAAGCTGGGTAAAATGGTCCAGATCTTGCTTCATT GGTC

***R10E8.8* promoter (∼1.6 kbp)**

F-R10E8.8p-Entry1- GGGGACAACTTTGTATAGAAAAGTTGCACATAAAATACGTTTTAGTAGCTGTCAG CAC

R-R10E8.8p-Entry1- GGGGACTGCTTTTTTGTACAAACTTGTTTTTGTCTGAAAATCGAACATTAAAAATA ACAGG

***elks-1* cDNA (∼2.5 kb)**

F-elks-1-cDNA-Entry2- GGGGACAAGTTTGTACAAAAAAGCAGGCTTTTCAGAAAAATGGCACCTGGTCCCGCACCATACAGC

R-elks-1-cDNA-Entry2- GGGGACCACTTTGTACAAGAAAGCTGGGTAGGCCCAAATTCCGTCAGCATCGTC GTG

*rab-3* promoter (∼1.2 kb) and *ges-1* promoter (∼3.4 kb) were used from de Bono lab plasmid collection.

#### Sequence of TurboID construct

GGAGGTGGTGGATCAGGCTCGGGAGGTCGAGGCTCAGGATCCGGTTCCGGCT

CCGGCTCTGGTTCCGGTTCGGGTTCCGGTTCTGGAAAGGATAACACCGTTCCA

CTTAAGCTTATCGCCCTTCTTGCCAACGGAGAATTCCACTCTGGAGAGCAACTT

GGAGAGACTCTTGGAATGTCCCGTGCTGCCATCAACAAGCATATCCAAACCCTT

CGTGATTGGGGAGTTGATGTTTTCACTGTTCCAGGAAAGgtaagtttaaacatatatatacta

actaaccctgattatttaaattttcagGGATACTCCCTTCCAGAGCCAATCCCACTTCTTAACGC

CAAGCAAATCCTTGGACAACTTGATGGAGGATCCGTCGCTGTCCTTCCAGTTGT

TGATTCCACCAACCAATACCTTCTTGACCGTATCGGAGAGCTTAAGTCTGGAGAC

GCCTGCATCGCTGAGTACCAACAAGCTGGACGCGGATCTCGCGGACGCAAGTG

GTTCTCCCCATTCGGAGCCAACCTTTACCTTTCTATGTTCTGGCGTCTTAAGCGT

GGACCAGCTGCTATCGGACTTGGACCAGTTATCGGAATCGTTATGGCTGAGGCC

CTTCGTAAGCTTGGAGCTGATAAGgtaagtttaaacagttcggtactaactaaccatacatatttaaattt

tcagGTTCGTGTTAAGTGGCCAAACGATCTTTACCTTCAAGACCGTAAGCTTGCTG

GAATCCTTGTCGAGCTTGCTGGAATCACCGGAGACGCCGCTCAAATCGTTATCG

GAGCTGGAATCAACGTTGCCATGCGTCGTGTTGAGGAGTCTGTTGTTAACCAAG

GATGGATCACTCTTCAAGAGGCTGGAATCAACCTTGATCGTAACACCCTTGCTG

CCACCCTTATCCGTGAGCTTCGTGCTGCCCTTGAGCTTTTCGAGCAAGAGGGAC

TTGCCCCATACCTTCCACGCTGGGAGAAGCTTGACAACTTCATCAACCGCCCAG

TTAAGCTTATCATCGGAGATAAGGAAATCTTCGGAATCTCTCGCGGAATCGACAA

GCAAGGAGCTCTTCTTCTTGAGCAAGATGGAGTCATTAAGCCATGGATGGGAGG

AGAGATTTCCCTTCGTTCCGCTGAGAAGGCCGGAGGAGAACAGAAGCTTATAAG

TGAGGAGGACCTGGGATCCGCTGGATCCGCTGCTGGATCCGGTGAGTTCATGG

TGTCGAAGGGAGAAGAGGATAACATGGCTTCACTCCCAGCTACACACGAACTCC

ACATCTTCGGATCGATCAACGGAGTGGATTTCGATATGGTCGGACAAGgtaagtttaa

acatatatatactaactaaccctgattatttaaattttcagGAACTGGAAACCCAAACGATGGATACGA

GGAACTCAACCTCAAGTCGACAAAGGGAGATCTGCAATTCTCGCCATGGATTCT

CGTGCCACACATCGGATACGGATTCCACCAATACCTCCCATACCCAGgtaagtttaaa

ctgagttctactaactaacgagtaatatttaaattttcagATGGAATGTCACCATTCCAAGCTGCCATG

GTGGATGGATCGGGATACCAAGTTCACCGAACAATGCAATTCGAGGATGGAGCC

TCGCTCACAGTGAACTACCGATACACATACGAGGGATCGCACATCAAGgtaagtttaa

acagttcggtactaactaaccatacatatttaaattttcagGGAGAGGCTCAAGTTAAGGGAACAGGA

TTCCCAGCTGATGGACCAGTGATGACAAACTCACTCACAGCTGCTGATTGGTGC

CGATCGAAAAAGACATACCCAAATGATAAGACAATCATCTCGACATTCAAGTGGT

CGTACACTACTGGAAACGGAAAGCGATACCGATCGACAGCCCGAACAACATACA

CATTCGCTAAGCCAATGGCCGCCAACTACCTCAAGgtaagtttaaacatgattttactaactaac

taatctgatttaaattttcagAATCAACCAATGTACGTGTTCCGAAAGACAGAACTCAAGCAC

TCAAAGACAGAGCTGAACTTCAAAGAGTGGCAAAAGGCCTTCACAGATGTGATG

GGAATGGATGAACTCTACAAGGACTACAAAGACCATGACGGTGATTATAAAGATC

ATGACATCGATTACAAGGATGACGATGACAAG

N/C terminus Gly-Ser rich linkers

CeTurboID

Introns

C-myc

mNeongreen

3XFLAG

### Light Microscopy

Confocal microscopy images of transgenic *C. elegans* expressing fluorescent proteins were acquired using a Leica (Wetzlar, Germany) SP8 inverted laser scanning confocal microscope with 10x 0.3 NA dry, 63x 1.2 NA water or 63x 1.2 NA oil-immersion objectives, using the LAS X software platform (Leica). The Z-project function in Image J (Rasband, W. S., ImageJ, U. S. National Institutes of Health, Bethesda, Maryland, USA, http://rsbweb.nih.gov/ij/) was used to obtain the figures used in the panels. Animals were mounted on 2% agarose pads and immobilized with 100 µM of sodium azide.

### Immunoblotting

Synchronized populations of *C. elegans* grown on *E. coli* MG1655 were harvested at L4 or young adult stage, washed three times in M9 buffer, incubated at room temperature (22°C) in M9 buffer supplemented with 1 mM biotin, and *E. coli* MG1655 for 2 hours. Two hours later the worms were washed three times in M9 buffer and flashfrozen after the M9 buffer was completely aspirated and 4X Bolt^TM^ LDS sample buffer supplemented with fresh DTT. The samples were then thawed, boiled for 10 minutes at 90°C, vortexed for 10 minutes, centrifuged for 30 minutes at 15000 rpm at 4°C and the supernatant collected. Proteins were transferred to a PVDF membrane (Thermofisher Scientific) following electrophoresis using Bolt 4-12% Bis-Tris Plus gels (Thermofisher Scientific). Membranes were blocked for 1 hour at room temperature with TBS-T buffer containing 5% BSA, and incubated for 2 hours at room temperature with HRP-conjugated or fluorescently-labeled streptavidin, or with HRP-conjugated antibodies. Membranes were then washed 3 times with TBS-T. The following antibodies or protein-HRP conjugates were used for this study: IRDye^®^ 800CW Streptavidin (1:7000 in TBS-T) (LI-COR Biosciences), Anti-FLAG M2- Peroxidase (1:5000 in TBS-T) (A8592 Sigma), anti-alpha tubulin-HRP (1:10000 in TBS-T) (DM1A Abcam ab40742), Streptavidin-HRP (1:5000 in TBS-T) (3999S Cell Signaling). Membranes were imaged using ChemiDoc the Imaging System (Models MP or XRS+, Bio-Rad).

### TurboID-based enzymatic protein labeling and extraction of biotinylated proteins from *C. elegans*

Gravid adult *C. elegans* were bleached and the eggs transferred to NGM plates seeded with *E. coli* MG1655 to obtain synchronized populations of worms. The animals were harvested at L4 or young adult stage, washed three times in M9 buffer, incubated at room temperature (22°C) in M9 buffer supplemented with 1 mM biotin, and *E. coli* MG1655 for 2 hours unless stated otherwise. Two hours later the worms were washed three times in M9 buffer and allowed to settle on ice after the last wash. After completely aspirating the M9 buffer, two volumes of RIPA buffer supplemented with 1 mM PMSF and cOmplete EDTA-free protease inhibitor cocktail (Roche Applied Science) was added to one volume of packed worms. The animals were again allowed to settle on ice and then added dropwise to liquid N_2_ to obtain frozen worm ‘popcorn’. A Spex 6875D cryogenic mill was used to grind frozen *C. elegans* to a fine powder which was then stored at -80°C. Worm powder was thawed by distributing it evenly along the length of a 50 ml falcon tube and rolling it on a tube roller at 4°C. After the sample was completely thawed, it was centrifuged (1000 rpm, 1 min, 4°C) to collect the sample at the bottom of the tube. SDS and DTT were added to the sample to final concentrations of 1% and 10mM respectively, from stock solutions of 20% SDS and 1M DTT. The tubes were gently inverted a few times and immediately incubated at 90°C for 5 minutes. After heat treatment, the samples were sonicated continuously for 1 minute twice, with brief cooling between the two sonication steps. Sonication used a probe sonicator microtip (MSE Soniprep 150 plus) and an amplitude setting of 16/max). The samples were cooled to room temperature following sonication and adjusted to 2M urea using a stock solution (8M urea, 1% SDS, 50 mM Tris-HCl, 150 mM NaCl). The samples were then centrifuged at 100000g for 45 minutes at 22°C, and the clear supernatant between the pellet and surface lipid layer transferred to a new tube.

Zeba^TM^ spin desalting columns (7K MWCO) (Thermofisher) were equilibrated 3 times with 5 ml RIPA buffer containing 1% SDS and 2M urea, freshly supplemented with protease inhibitors (Roche cOmplete EDTA-free protease inhibitor cocktail 1 tablet/25 ml; PMSF 1 mM) by centrifugation at 1000g for 5 minutes (or until the buffer was completely eluted from resin). Around 4 ml of clarified sample was then loaded onto the equilibrated spin column and desalted by centrifugation at 1000g for 5 minutes (or until the sample was completely eluted from resin) to remove free biotin. The desalting step was repeated once more using freshly equilibrated columns. Protein concentration in the samples was measured using Pierce^TM^ 660nm protein assay reagent supplemented with Ionic Detergent Compatibility Reagent (IDCR) (ThermoFisher Scientific).

### Biotinylated protein pulldown and elution

Dynabeads^TM^ MyOne^TM^ streptavidin C1 (Invitrogen, 2 ml bead slurry per 90 mg of *C. elegans* total protein lysate) were equilibrated in RIPA buffer by briefly incubating them three times in the buffer and using magnetic separation to retain beads while discarding buffer (note: we were able to reduce the amount of protein lysate to 5-10 mg in subsequent experiments without compromising the mass spectrometry data). Desalted, clarified samples were combined with Dynabeads in a 50 ml falcon tube and incubated overnight in a tube roller at room temperature. Beads were magnetically separated using a neodymium magnet taped to the tube and incubated on a rocking platform. Unbound lysate was removed and the Dynabeads transferred to 2 ml LoBind protein tubes (Eppendorf). Beads were washed five times with 2% SDS wash buffer (150 mM NaCl, 1 mM EDTA, 2% SDS, 50 mM Tris-HCl, pH 7.4) and five times with 1M KCl wash buffer; beads were transferred to new tubes after each wash.

Elution sample buffer was prepared by dissolving free biotin (Sigma) to saturation in sample buffer (NuPAGE LDS sample buffer 4X) containing reducing agent (NuPAGE sample reducing agent 10X). Elution sample buffer was centrifuged at 13000 rpm for 5 minutes to remove undissolved biotin and the elution sample buffer saturated with dissolved biotin was transferred to a new tube. 100 µl of elution buffer was applied to Dynabeads, vortexed gently, heated for 5 minutes at 90°C, before the vortexing and heating steps were repeated. Dynabeads were separated magnetically and elution buffer containing biotinylated proteins was transferred to a new 1.5 ml LoBind protein tube (Eppendorf). 70 µl of sample was electrophoresed on a NuPAGE^TM^ 4-12% Bis-Tris protein gel (Invitrogen), which was then stained with InstantBlue (Expedeon) for visualization. The gel was sliced into 20 fractions, and sent for mass spectrometry analysis.

### Mass spectrometry

Polyacrylamide gel slices (1-2 mm) containing the purified proteins were prepared for mass spectrometric analysis using the Janus liquid handling system (PerkinElmer, UK). Briefly, the excised protein gel pieces were placed in a well of a 96-well microtitre plate, destained with 50% v/v acetonitrile and 50 mM ammonium bicarbonate, reduced with 10 mM DTT, and alkylated with 55 mM iodoacetamide. After alkylation, proteins were digested with 6 ng/µL Trypsin (Promega, UK) overnight at 37 °C. The resulting peptides were extracted in 2% v/v formic acid, 2% v/v acetonitrile. The digest was analysed by nano-scale capillary LC-MS/MS using an Ultimate U3000 HPLC (ThermoScientific Dionex, San Jose, USA) to deliver a flow of approximately 300 nL/min. A C18 Acclaim PepMap100 5 µm, 100 µm x 20 mm nanoViper (ThermoScientific Dionex, San Jose, USA), trapped the peptides prior to separation on an EASY-Spray PepMap RSLC 2 µm, 100 Å, 75 µm x 250 mm nanoViper column (ThermoScientific Dionex, San Jose, USA). Peptides were eluted using a 60-minute gradient of acetonitrile (2% to 80%). The analytical column outlet was directly interfaced via a nano-flow electrospray ionisation source with a hybrid quadrupole orbitrap mass spectrometer (Q-Exactive Orbitrap, ThermoScientific, San Jose, USA). Data collection was performed in data-dependent acquisition (DDA) mode with an r = 70,000 (@ *m*/*z* 200) full MS scan from *m*/*z* 380–1600 with a target AGC value of 1e6 ions followed by 15 MS/MS scans at r = 17,500 (@ *m*/*z* 200) at a target AGC value of 1e5 ions. MS/MS scans were collected using a threshold energy of 27 for higher energy collisional dissociation (HCD) and a 30 s dynamic exclusion was employed to increase depth of coverage. LC-MS/MS data were then searched against a protein database (UniProt KB) using the Mascot search engine programme (Matrix Science, UK) (*42*). Database search parameters were set with a precursor tolerance of 10 ppm and a fragment ion mass tolerance of 0.8 Da. One missed enzyme cleavage was allowed and variable modifications for oxidized methionine and N-terminal acetylation were included. Carbamidomethylation of cysteine was specified as a fixed modification. MS/MS data were validated using the Scaffold programme (Proteome Software Inc., USA) (*43*). All data were additionally interrogated manually.

## Acknowledgements

We thank de Bono lab members for helpful comments on the manuscript, IST Austria and University of Vienna Mass Spec Facilities for invaluable discussions and comments for the optimization of mass spec analyses of worm samples. The biotin auxotropic *E. coli* strain MG1655bioB:kan was gift from John Cronan (University of Illinois) and was kindly sent to us by Jessica Feldman and Ariana Sanchez (Stanford University). dg398 pEntryslot2_mNeongreen::3XFLAG::stop and dg397 pEntryslot3_mNeongreen::3XFLAG::stop::unc-54 3′UTR entry vector were kindly shared by Dr. Dominique Glauser (University of Fribourg). Codon optimized mScarlet vector was a generous gift from Dr. Manuel Zimmer (University of Vienna). This work was supported by an Advanced ERC Grant (269058 ACMO) and a Wellcome Investigator Award (209504/Z/17/Z) to MdB and an ISTplus Fellowship to MA (Marie Sklodowska-Curie agreement No 754411).

## Author Contributions

MA, SB and MdB conceived experiments; MA, SB and SMF performed experiments; MA, SB and MdB analysed data; FB and MS performed mass spectrometry analysis and SB and AN processed mass spectrometry data; MA and MdB wrote the manuscript.

## Supplementary Materials

**Fig. S1.**
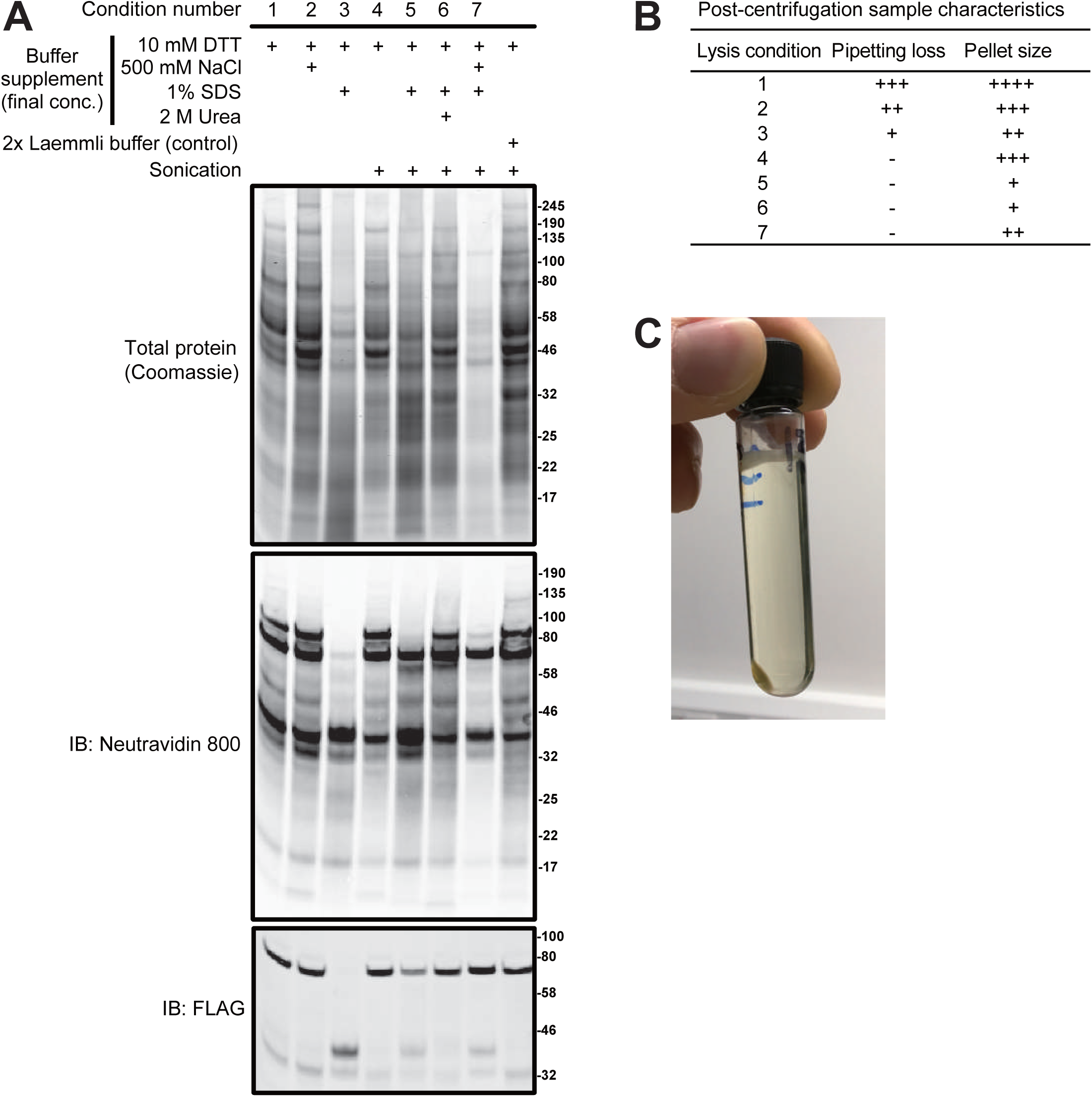
Optimizing extraction and solubilization of *C. elegans* proteins. **(A**) Different extraction protocols compared by visualizing total protein (Coomassie-stained gel, top), biotinylated protein (Western blot probed with Neutravidin 800), and the FLAG-tagged TurboID fusion (Western blot probed with anti-FLAG antibody). Tissue samples were from worms expressing pan-neuronal free TurboID (*rab-3p::TbID::mNG*). Protocol 5 best extracted and solubilized proteins. (**B**), Post-centrifugation pellet sizes obtained with the 7 extraction conditions in (A). (**C**), Typical pellet appearance following clarification by ultracentrifugation following selected extraction method. showing >=95% solubilization by volume. Tube capacity in the image in 10.4 ml

**Fig. S2.**
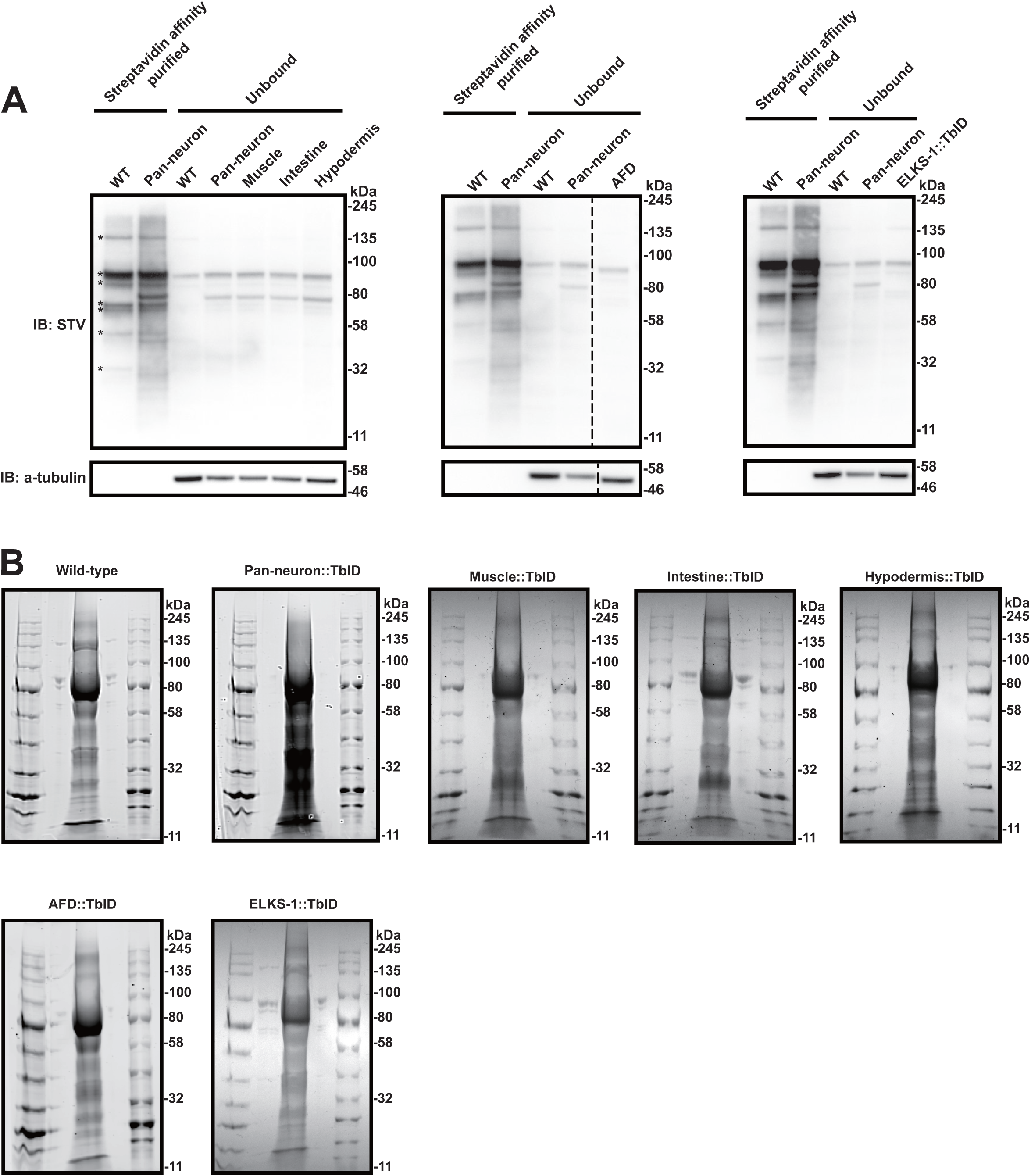
Testing the efficiency of streptavidin affinity purification of biotinylated proteins. (**A**) Western blot analysis of bound and unbound biotinylated proteins after streptavidin-based affinity purification of samples. (**B**) Proteins isolated from various samples and affinity purified using streptavidin beads, separated using 4-12% Bis-Tris gels and stained by Coomassie. Also shown are molecular weight markers. Slices from these gels were sent for mass spectrometry.

**Fig. S3.**
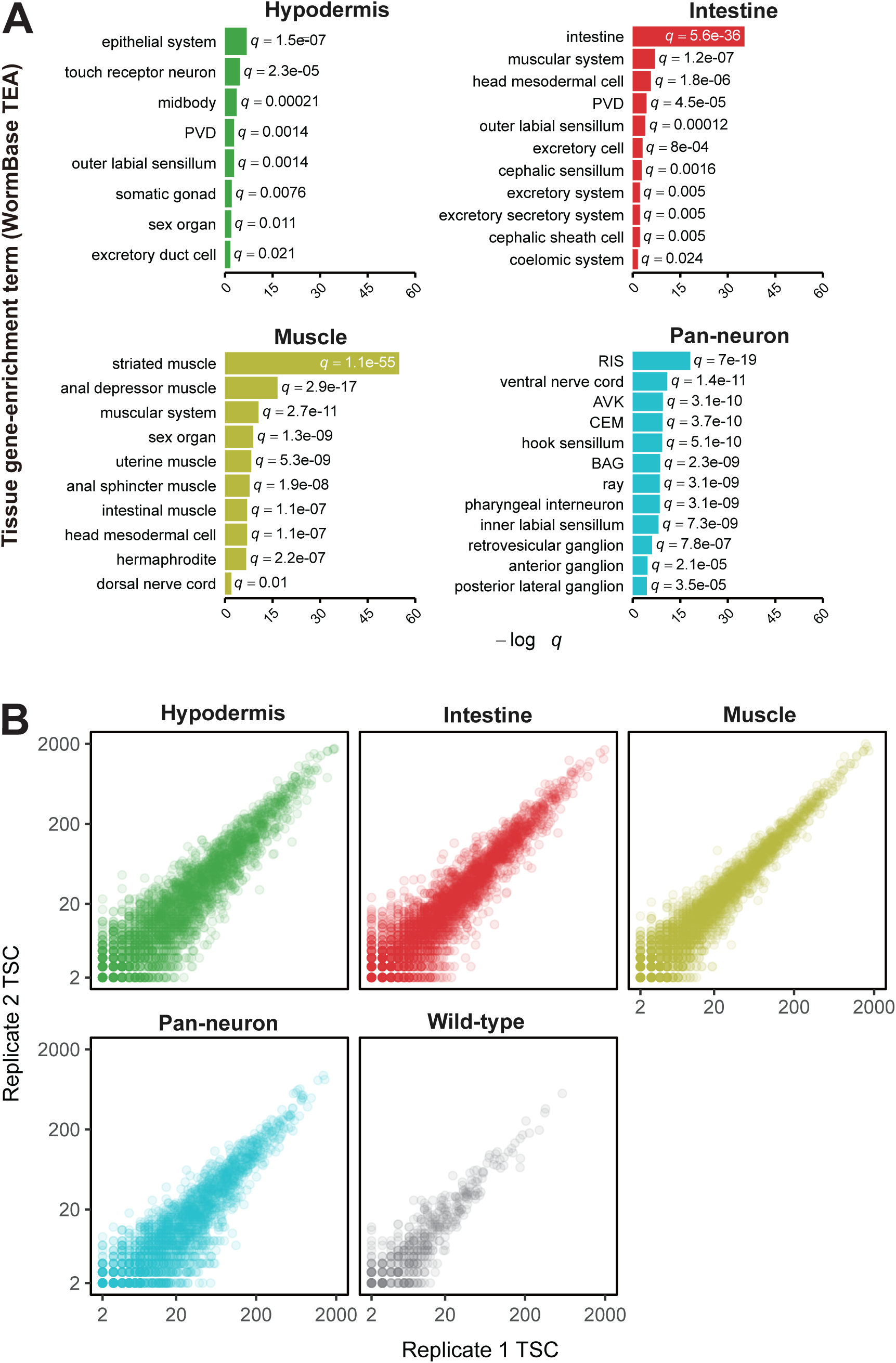
Tissue enrichment analysis of unique protein hits identified by mass spectrometry analysis of four major *C. elegans* tissues. (**A**) Top-ranked Wormbase tissue-enrichment analysis (TEA) terms associated with tissue-enriched protein hits detected across indicated samples. (**B**) Replicate 1 vs replicate 2 total spectral counts for protein hits from indicated samples. Points correspond to individual proteins identified.

**Fig. S4.**
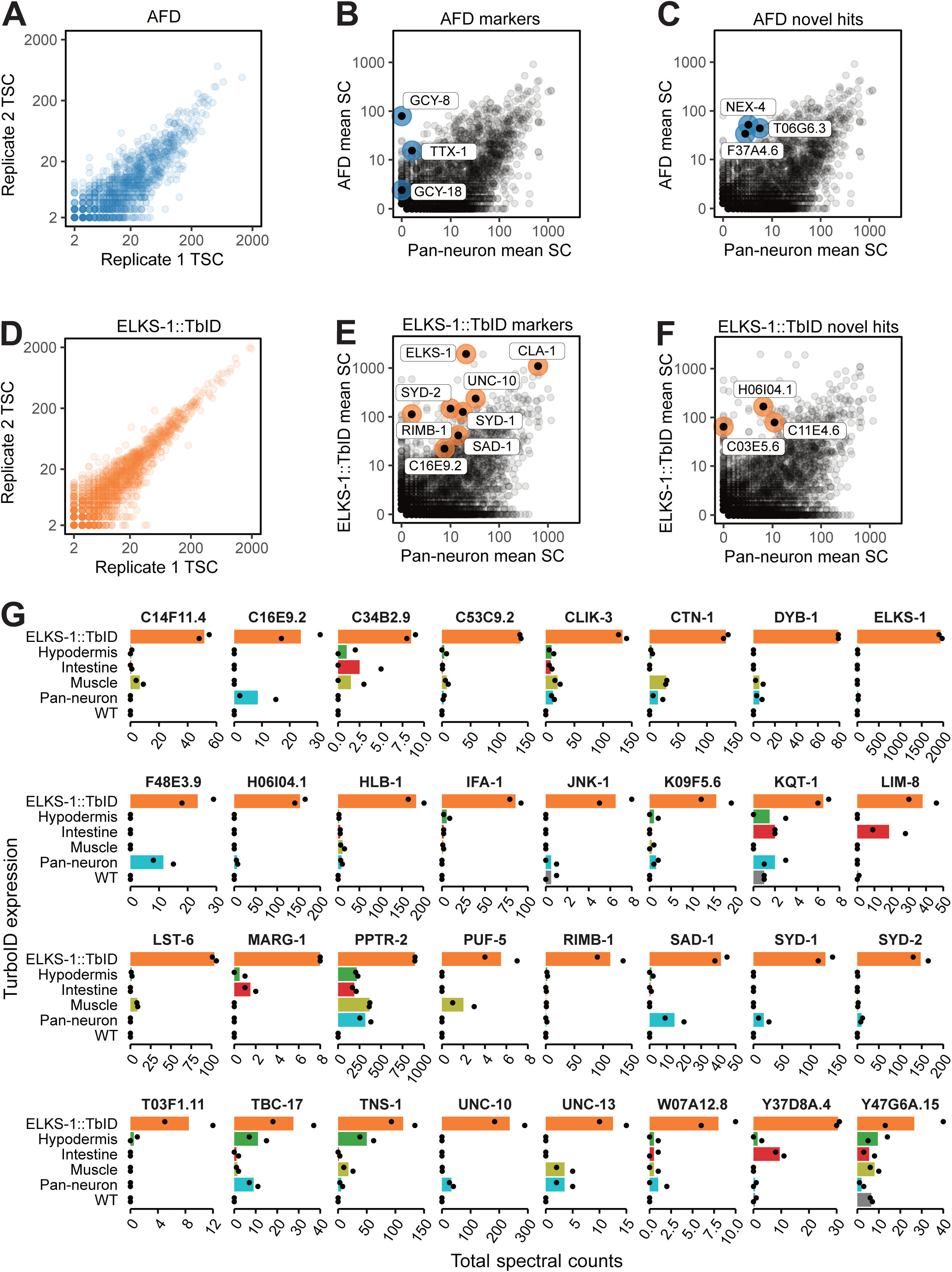
Spectral counts analysis of proteins across replicate samples of AFD-specific TurboID and ELKS-1::TurboID. (**A**) Replicate 1 vs replicate 2 total protein counts for protein hits. Points correspond to individual proteins identified in AFD neurons. (**B** and **C**) Comparison of protein mean spectral counts between AFD and pan-neuronal free TurboID samples. Known marker proteins (B) and novel hits (C) are highligted. (**D**) Replicate 1 vs replicate 2 total protein counts for protein hits. Points correspond to individual proteins identified in ELKS-1::TurboID. (**E** and **F**) Comparison of protein mean spectral counts between ELKS-1::TurboID and pan-neuronal free TurboID samples. Known marker proteins (E) and novel hits (F) are highlighted. (**G**) Proteins enriched two fold or more in ELKS-1::TurboID samples compared to other samples, and represented by a mean of at least 5 spectral counts.

## Supplemental table legends

Table S1: Total spectral counts of protein hits identified in tissues, AFD and ELKS-1::TbID samples.

